# Engineering recombination between diverged yeast species reveals genetic incompatibilities

**DOI:** 10.1101/755165

**Authors:** G. Ozan Bozdag, Jasmine Ono, Jai A. Denton, Emre Karakoc, Neil Hunter, Jun-Yi Leu, Duncan Greig

**Author notes:** Corresponding author: Jasmine Ono. Authors contributed equally (G. Ozan Bozdag, Jasmine Ono).

## Abstract

The major cause of the sterility of F1 hybrids formed between *Saccharomyces cerevisiae* and *Saccharomyces paradoxus* is anti-recombination. The failure of homologous chromosomes from the different species to recombine causes them to mis-segregate, resulting in aneuploid gametes, most of which are inviable. These effects of anti-recombination have previously impeded the search for other forms of incompatibility, such as negative genetic interactions (Bateson-Dobzhoansky-Muller incompatibilities). By suppressing the meiotic expression of *MSH2* and *SGS1*, we could increase recombination and improve hybrid fertility seventy-fold. This allowed us to recover meiotic tetrads in which all four gametes were viable, ensuring that segregation had occurred properly to produce perfectly haploid, not aneuploid, recombinant hybrid gametes. We sequenced the genomes of 84 such tetrads, and discovered that some combinations of alleles from different species were significantly under-represented, indicating that there are incompatible genes contributing to reproductive isolation.

## Introduction

Species are formed and maintained by the restriction of gene flow between diverging populations. Barriers to gene flow can be physical, such as geographic distance, or they can be properties of the species themselves. Here, we focus on one such barrier to gene flow, hybrid sterility. Hybrid sterility is a form of post-zygotic reproductive isolation, meaning that it acts after diverging populations have already mated and produced a hybrid zygote. Hybrid sterility can be caused by a variety of mechanisms that can generally be classified into incompatibilities between diverged chromosomes (such as large-scale chromosomal rearrangements; Rieseberg and Willis, 2007, and anti-recombination) and incompatibilities between individual genes from the diverging populations (Presgraves, 2010). There is particular interest in the latter class of genic incompatibilities, which are often referred to as “Bateson-Dobzhansky-Muller incompatibilities” (BDMIs) or “speciation genes” (Orr, 1996). As we don’t know whether these incompatibilities themselves are the cause of speciation or have developed post-speciation, we will refer to them as BDMIs throughout.

BDMIs represent a case where alleles (at two or more loci) that have evolved to work well together within a species perform poorly when combined in a hybrid individual with alleles from another species, whose alleles have evolved independently (Coyne and Orr, 2004). Since BDMIs offer a universal mechanism for speciation, they have been studied intensely, both theoretically and empirically, yet only a handful have been discovered and characterized at the molecular level (reviewed in Presgraves, 2010; Rieseberg and Blackman, 2010; Maheshwari and Barbash, 2011; Nosil and Schluter, 2011). Understanding the molecular mechanisms underlying additional BDMIs will allow us to address general questions about reproductive isolation, such as the number of BDMIs typically involved and their effect sizes, whether the same genes or types of genes are involved in different cases, what types or locations of mutations are most likely to cause incompatibility, and whether BDMIs evolve by selection or drift (Nosil and Schluter, 2011).

Yeast are a great system in which to molecularly characterize such interactions because the genomic data, molecular tools and genetic tractability of the model yeast *Saccharomyces cerevisiae* are unsurpassed by any other model eukaryote (Botstein and Fink, 2011). F1 hybrids between *S. cerevisiae* and its closest relative *Saccharomyces paradoxus* have greatly reduced sexual fertility compared to non-hybrids. Haploid gametes from the two different species can fuse to form diploid F1 hybrids that grow normally by mitosis, but only about 1% of the gametes (which are produced as spores) formed via meiosis are viable (able to germinate and grow into colonies) (Hunter et al., 1996). In contrast, nearly all the spores produced by non-hybrids are viable. The two species do not differ by substantial chromosomal rearrangements that might account for this sterility (Fischer et al., 2000; Kellis et al., 2003). Instead, a form of chromosomal incompatibility known as anti-recombination is thought to be the cause. The two species’ genomes are so diverged in sequence (about 12% of nucleotide positions differ; Rogers et al., 2018) that homologous recombination is suppressed, and meiotic crossing over is greatly reduced. Because crossovers are important for chromosome segregation during meiosis, efficient segregation is impaired, and gametes are killed because they lack one or more essential chromosomes or, potentially, because they carry extra chromosomes. Consistent with this, the 1% viable gametes produced from hybrid meioses are aneuploid, carrying additional chromosomes, and very few chromosomes are recombinant (Hunter et al., 1996; Kao et al., 2010).

In principle, chromosome mis-segregation alone is capable of explaining yeast hybrid sterility without invoking any role for BDMIs. We recently quantified the precise rates at which each chromosome segregates in F1 hybrids (Rogers et al., 2018). The average rate of correct distribution for each chromosome in hybrids formed between *S. cerevisiae* and *S. paradoxus* is 59.7%, so we expect only 0.03% of gametes to receive exactly one copy of each chromosome (0.597 for each chromosome, raised to the power of 16 to account for all sixteen chromosomes). However, gametes carrying more than one copy of a chromosome can also be viable, as shown by the high rates of aneuploidy detected in viable hybrid gametes. In the 40.3% of hybrid meioses in which a chromosome does not segregate properly, half of the resulting spores (20.15%) will receive two copies of the chromosome and might therefore be viable, whilst the remaining 20.15% will receive no copies and will certainly be inviable. Therefore 2.7% of gametes (0.597 plus 0.2015, raised to the power of 16) will receive *at least* one copy of each essential chromosome, and could be viable, depending on the effect of the additional chromosomes that they carry (Boynton et al., 2018). Thus chromosome mis-segregation due to anti-recombination accounts for at least 97.3%, and potentially all, of the observed hybrid sterility. However, there is little direct evidence that extra chromosomes contribute to spore inviability (Rogers et al., 2018), so the smaller figure is more likely, leaving open the possibility that some hybrid spores are killed because of incompatible interactions between genes of one species and those of the other (BDMIs).

To date, no such BDMIs have been detected in yeast. BDMIs have been detected between mitochondrial genes from one yeast species and nuclear genes from another (Lee et al., 2008; Chou et al., 2010; also see Xu and He, 2011), but these act earlier by reducing F1 mitotic viability and preventing F1 meiosis from even occurring, not by causing inviability of the gametes produced by hybrid meiosis. We have previously shown that most *S. paradoxus* chromosomes can successfully replace their homologues in *S. cerevisiae* haploid gametes when substituted one at a time, indicating that they do not contain always-lethal incompatibilities (Greig, 2007). But this method would not detect weaker BDMIs that kill only sometimes (incomplete penetrance), or that have a cumulative effect with other BDMIs on other chromosomes. A possible way to detect such BDMIs is to genotype the surviving gametes from hybrid meioses and test whether some combinations of alleles from different species at different loci are statistically under-represented. The explanation for such under-representation would be that they are incompatible and cause gamete inviability. This method has been modelled by Li et al. (2013), and has been implemented by Kao et al. (2010). Whilst the distribution of genotypes differed significantly from what was expected by chance, the additional aneuploid chromosomes carried by the genotyped gametes confounded analysis to an extent that the effective sample size was too low to identify individual pairs of incompatible loci (Kao et al., 2010).

In order to identify BDMIs involved in hybrid spore inviability, it is therefore necessary to overcome the primary effect of anti-recombination, in order to produce haploid spores without additional aneuploid chromosomes for genotyping. Hunter et al. (1996) previously showed that knocking out genes involved in monitoring the fidelity of recombination increases both the rate of recombination and the proportion of viable gametes produced by hybrid meioses. By deleting the mismatch repair gene *MSH2*, they increased crossing over in hybrids on average 13-fold, resulting in a nearly 9-fold increase in hybrid spore viability. Kao et al. (2010) therefore used msh2? knock-out mutants in their search for BDMIs, but the improvement in chromosome segregation was insufficient to relieve the extensive aneuploidy of the hybrid gametes. Here we employed two additional tools in order to produce perfectly euploid hybrid gametes for genotyping. First, we repressed the expression of both *MSH2* and a second anti-recombination factor, DNA-helicase *SGS1*, specifically in meiosis, thereby retaining their normal function during mitosis, which reduces the mutagenic effects of knocking them out entirely. Secondly, we dissected hybrid gametes out of their meiotic tetrads and genotyped only those that came from tetrads in which all four spores were viable. Our sample therefore excluded not only those gametes containing lethal combinations of the parent species’ alleles, but also aneuploid gametes, since any chromosome mis-segregation will kill some of the gametes in a tetrad.

We sequenced all 336 haploid gametes from 84 F1 hybrid meioses and tested statistically for pairs of alleles for which parental combinations were over-represented. We were able to map four broad pairs of genomic regions that show evidence of incompatibility. Thus, for the first time, we find evidence of naturally-occurring nuclear BDMIs causing sterility of hybrids between two species of yeast.

## Results

### Restoration of hybrid fertility

We constructed strains of *S. cerevisiae* and *S. paradoxus* in which the native promoters of *MSH2* and *SGS1* were replaced with the *CLB2* promoter, which is specifically repressed during meiosis (Grandin and Reed, 1993; Lee and Amon, 2003) (Table 1, Supplementary File 1, Supplementary File 2). *MSH2* and *SGS1* are both implicated in the anti-recombination process (Chakraborty and Alani, 2016). By maintaining expression of these genes in mitosis, we can avoid any unwanted effects such as an increased recessive-lethal mutation rate, which would actually reduce fertility (Hunter et al., 1996). In a previous study, we found that suppressing meiotic expression of *SGS1* alone improved the rate of correct segregation by almost half in hybrid meioses (Rogers et al., 2018). Here, we find that spore viability is also dramatically improved. Suppression of *SGS1* alone increased hybrid spore viability from 0.46% to 20.8%; and in combination with suppression of *MSH2*, spore viability was further improved to 32.6% (Figure 1, Source Data 1). Significantly more of the double mutant spores were viable than in the wild-type hybrid (chi-squared contingency test: X^2^ = 479.91, df = 1, p-value < 2.2×10^−16^).

**Table 1:** List of strains used in this study. For a complete list, see Supplementary File 2.

| YDG | Strain name | Original strain name |  |
| --- | --- | --- | --- |
| 391 | NCYC 3708 | N17 | ho::HYGMX @ ura3::KanMX |
| 542 | NCYC 3583 | W303 | ho::HYGMX a ura3::KanMX ade2-1 |
| 832 | NHY 2039 | | ho::hisG a ura3(ΔSma-Pst) HIS4::LEU2-(BamHI; +ori) leu2::hisG pCLB2-3HA-SGS1::kanMX4 |
| 853 | YDG391 x YDG542 | N17 x W303 | ho::HYGMX/ho::HYGMX a/@ ura3::KanMX/ura3::KanMX ade2-1/ADE2 |
| 866 |  | W303 | ho a ura his3 leu2::NAT trp ade-2 can1r pCLB2-3HA-SGS1::kanMX4 |
| 905 |  | N17 | ho @ ura3 cyh2r pCLB2-3HA-SGS1::kanMX4 |
| 912 | YDG866 x YDG905 | W303 x N17 | ho/ho a/@ ura/ura3 his3/HIS3 leu2::NAT/LEU2 trp/TRP ade-2/ADE can1r/CAN1 CYH2/cyh2r pCLB2-3HA-SGS1::kanMX4/pCLB2-3HA-SGS1::kanMX4 |
| 959 |  | N17 | ho a lys2 cyh2r pCLB2-3HA-MSH2::kanMX4 |
| 960 | YSC1059 | W303 | ho @ ura3-52 his3-11 leu2-3,112 trp1Δ2 ade2-1 can1-100 pCLB2-3HA-MSH2::kanMX4 |
| 964 | YDG959 x YDG960 | N17 x W303 | ho/ho a/@ URA3/ura3-52 lys2/LYS2 HIS3/his3-11 LEU2/leu2-3,112 TRP1/trp1Δ2 ADE2/ade2-1 CAN1/can1-100 cyh2r/CYH2 pCLB2-3HA-MSH2::kanMX4/pCLB2-3HA-MSH2::kanMX4 |
| 967 |  | N17 | ho a cyh2r pCLB2-3HA-SGS1::kanMX4 pCLB2-3HA-MSH2::kanMX4 |
| 968 |  | N17 | ho @ cyh2r pCLB2-3HA-SGS1::kanMX4 pCLB2-3HA-MSH2::kanMX4 |
| 969 |  | W303 | ho a ura3 his3 leu2::NAT trp1 ade-2 can1r pCLB2-3HA-SGS1::kanMX4 pCLB2-3HA-MSH2::kanMX4 |
| 970 |  | W303 | ho @ ura3 his3 leu2::NAT trp1 ade-2 can1r pCLB2-3HA-SGS1::kanMX4 pCLB2-3HA-MSH2::kanMX4 |
| 982 | YDG968 x YDG969 | N17 x W303 | ho/ho @/a URA3/ura3 HIS3/his3 LEU2/leu2::NAT TRP1/trp1 ADE/ade-2 CAN1/can1r cyh2r/CYH2 pCLB2-3HA-SGS1::kanMX4/pCLB2-3HA-SGS1::kanMX4 pCLB2-3HA-MSH2::kanMX4/pCLB2-3HA-MSH2::kanMX4 |

**Figure 1:**
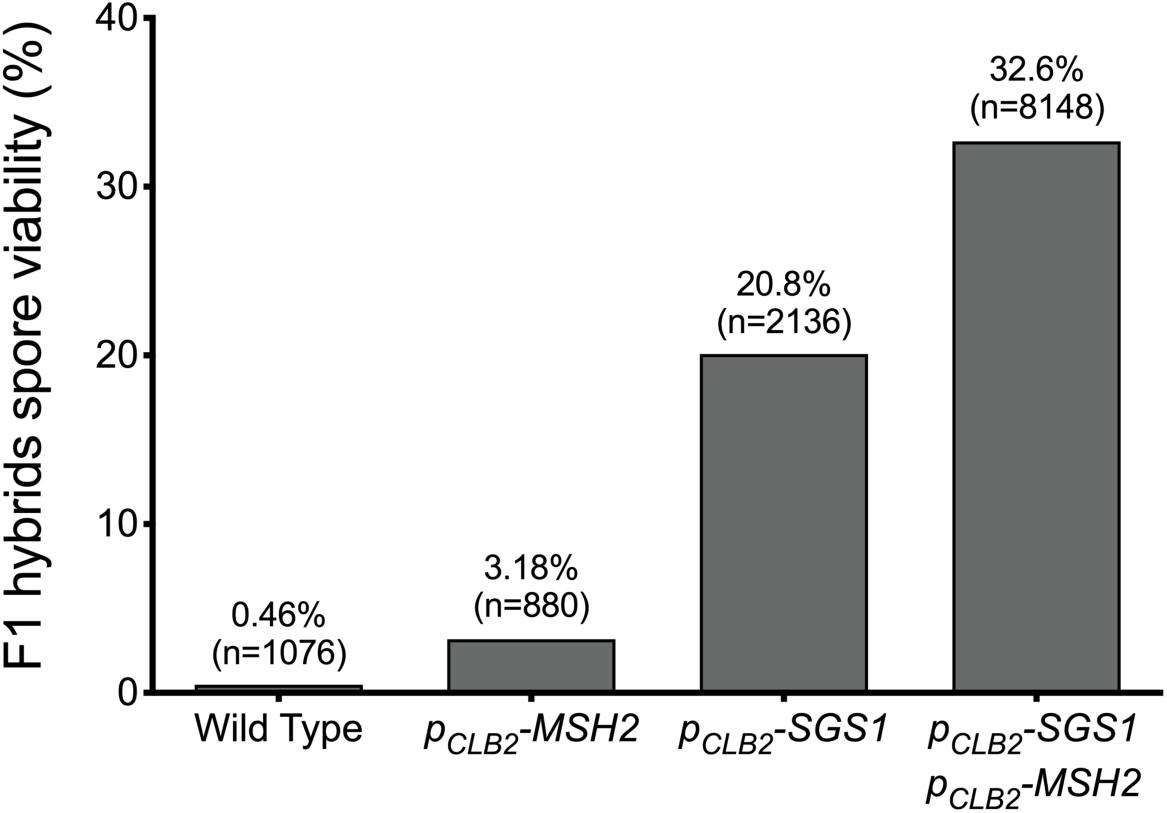
Restoration of hybrid fertility by meiotic repression of *MSH2* and *SGS1*. Percentages are spore viabilities of the indicated hybrid strains. In the *P*_*CLB2*_*-MSH2 P*_*CLB2*_*-SGS1* strain, a significant 32.14% increase in spore viability was observed (double mutant when compared with the wild type: X^2^ = 479.91, df = 1, p-value < 2.2×10^−16^). Numbers in parentheses indicate the total number of dissected spores checked for viability. Full data, including other strains, can be found in Source Data 1.

The restoration of hybrid fertility vastly increased the production of hybrid tetrads in which all four spores were viable, which were specifically selected for genotyping and further analysis. All spores from such tetrads are necessarily euploid, as mis-segregation of even a single chromosome would result in at least one dead spore (lacking that chromosome). By analyzing only euploid spores, we ensured that recessive BDMIs were not masked by aneuploidy.

### Evidence for hybrid incompatibility

A fertility-reducing BDMI between a pair of loci would result in fewer gametes containing hybrid combinations of alleles at these loci. Reasoning that we could not map such loci at a resolution higher than the linkage groups produced by the crossovers that occurred within the 84 tetrads in our sample, we divided the chromosomes into 1208 segments defined by all of the recombination breakpoints produced by our genotyping procedure (see Supplementary File 3). Treating each of these segments as a putative incompatibility locus, we tested every segment against every other segment, excluding those on the same chromosome, using two-by-two contingency tables in the manner described by Li et al. (2013). Those on the same chromosome cannot be tested because physical linkage cannot be distinguished from linkage due to interaction. We calculated the odds ratio (OR) for each pair by dividing the product of the numbers of the two parental genotypes observed by that of the two hybrid types:

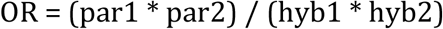

In addition, we calculated the 99% confidence interval (CI) for the odds ratios. An odds ratio of 1 indicated that the parental and hybrid types were present in equal frequencies. An odds ratio greater than 1 would be observed for a bias towards parental types; and an odds ratio less than 1 would indicate that hybrid types were preferentially observed. We found all pairwise comparisons for which the calculated 99% CI did not encompass the value of 1 (lower bound of CI > 1 or upper bound of CI < 1). 1.9% of all comparisons (13,082/676,294) had CIs that indicated a hybrid preference and 2.6% (17,492/676,294) had CIs that indicated a parental preference. Parental types were not only over-represented more often, but were also more highly favoured. Parental types were over-represented by 3/4 in 190 cases (parent/hybrid ratio ≧ 1.75) while hybrid types were over-represented by 3/4 in only 22 cases (hybrid/parent ratio ≧ 1.75).

Individual significant interactions were determined as described in Li et al. (2013) and in the Methods. Briefly, a null distribution of top ORs was produced by randomly re-sampling the observed data 100 times (see Source Code 1). The 5th largest OR from this set of the top 100 ORs was used as the critical value from which we judged significance. All observed pairs with a higher OR than the critical value (3.41) were deemed significant (Supplementary File 4 and Source Data 2). Blocks were then formed from these significant pairs by grouping nearby segments that interacted with other nearby segments (see Methods for details). Of note, region B interacts with a segment adjacent to those interacting with region F (Figure 2). We collapsed both interacting regions into one (region D) as the two interactions may involve a single underlying gene in the broader area. In this way, we found four putative BDMIs involving six regions of the genome: between regions A and E (two significant interactions - highest OR = 4 or 2-fold over-representation of parental combinations), regions B and D (one significant interaction - highest OR = 3.60 or 1.90-fold difference), regions C and E (eight significant interactions - highest OR = 4.71 or 2.17-fold difference) and regions D and F (32 significant interactions - highest OR = 4 or 2-fold difference) (Table 2, Figure 2). All four interactions involve only four chromosomes, with multiple independently significant interactions mapping to the same regions.

**Figure 2:**
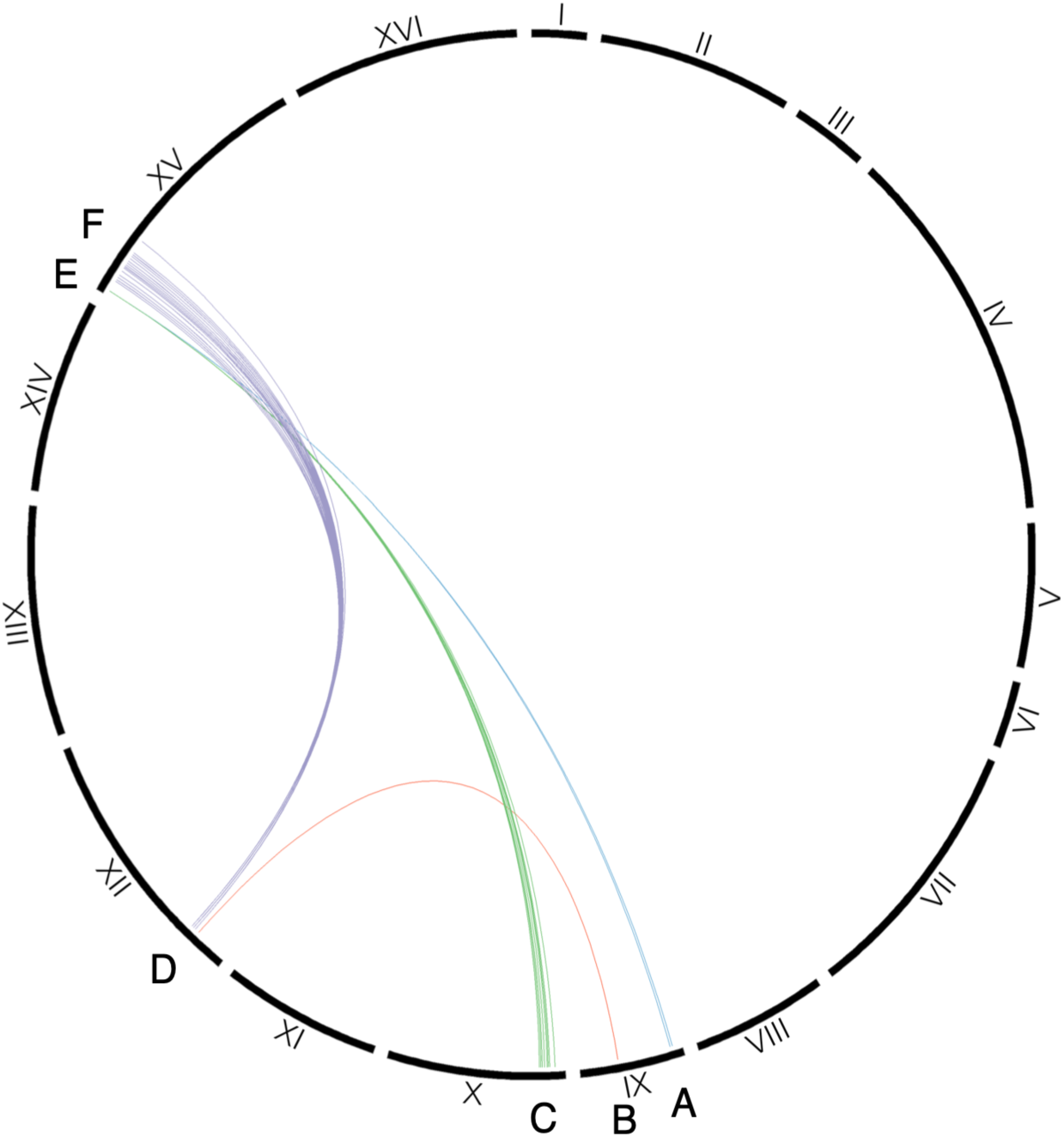
Four putative BDMIs mapped to six genomic regions. Here, the chromosomes are displayed in a circle and each significant pairwise interaction is indicated by a line linking the involved segments. The interactions were grouped by nearby segments, forming six interacting regions and four putative BDMIs. Interactions contributing to different putative BDMIs are coloured differently. For the code used to determine significance, see Source Code 1. Source Data 2 contains all significant interactions.

**Table 2:** The six genomic regions involved in putative BDMIs. Region E interacts with both A and C, and region D interacts with both B and F. The ranges and number of genes are based on SGRP sequencing added 10/10/08 (Liti et al., 2009; Bergström et al., 2014) and can be determined using Source Data 2 in conjunction with Supplementary File 3.

| Region | Chr | Genomic range (W303) | Number of genes (W303) | Genomic range (N17) | Number of genes (N17) |
| --- | --- | --- | --- | --- | --- |
| A | IX | 32385-52752 | 13 | 11474-28361 | 11 |
| B | IX | 275518-276372 | 1 | 248868-249722 | 1 |
| C | X | 25663-113149 | 55 | 2529-101136 | 54 |
| D | XII | 138633-196103 | 30 | 125307-183140 | 30 |
| E | XV | 16405-26386 | 5 | 2188-14304 | 5 |
| F | XV | 63347-279621 | 121 | 51621-263564 | 117 |

## Discussion

### Anti-recombination as a barrier between species

By reducing the expression of just two genes, *SGS1* and *MSH2*, during meiosis we were able to rescue the fertility of a sterile hybrid between *S. cerevisiae* and *S. paradoxus*, increasing its ability to produce viable gametes 70-fold, from 0.46% to 32.6%. The fertility of our rescued inter-species hybrid was around one third that of its non-hybrid parents, which is about the same as intra-species crosses formed between diverged populations of a single species, *S. paradoxus* (e.g., Greig et al., 2003; Charron et al., 2014). These anti-recombination genes thus determine most of the hybrid sterility barrier between *S. cerevisiae* and *S. paradoxus*.

The hybrid sterility effects of *SGS1* and *MSH2* are not caused by incompatibilities between their alleles from the different species, but rather by their effects on the physical interaction between whole chromosomes from the different species. In non-hybrids, the Sgs1 and Msh2 proteins act to physically impede the formation and stabilization of heteroduplex DNA formed during recombination between mismatched DNA sequences. This activity helps to maintain genome integrity by permitting recombination at allelic positions between matching homologous chromosomes, but preventing ectopic (non-allelic) recombination between non-homologous chromosomes and dispersed repeats, which would cause rearrangements. However, the genomes of *S. cerevisiae* and *S. paradoxus* differ by 12% at allelic positions across the genome (Rogers et al., 2018), so there are enough mismatches to globally reduce meiotic recombination between homologous chromosomes in hybrids. Kao et al. (2010) found that viable hybrid spores had only 2.7 crossovers per meiosis. By inferring crossover rates in dead spores, Rogers et al. (2018) measured an overall rate of just one crossover per hybrid meiosis, much lower than the normal rate of about 90 crossovers that occur in a non-hybrid *S. cerevisiae* meiosis (Martini et al., 2006). Thus most of the sixteen pairs of chromosomes in a hybrid lack any meiotic crossovers, leading to aneuploidy and inviability in the spores.

Manipulating the expression of *SGS1* has previously been shown to greatly improve meiotic segregation of chromosomes from different species, both in partial hybrids, in which only one chromosome comes from another species, and in full hybrids as we used here. Amin et al. (2010) found that meiotic non-disjunction of a single chromosome III from *S. paradoxus* in an otherwise *S. cerevisiae* background fell 2.5-fold, from 11.5% to 4.6% per meiosis, when *SGS1* was repressed during meiosis using the *CLB2* promotor. Rogers et al. (2018) found that the non-disjunction rate in full hybrids was much higher than in partial hybrids, averaging 40.3% per chromosome per meiosis. Nevertheless, repressing *SGS1* expression also improved segregation in the full hybrids by between 2-fold and 3.2-fold, depending on the chromosome. Here we showed that this improvement in segregation is sufficient to greatly improve fertility, confirming that anti-recombination comprises the major component of the species barrier.

The much larger effect of repressing *SGS1* expression on hybrid viability, relative to that of repressing *MSH2*, could be explained by Sgs1 having several effects on homolog interactions and meiotic recombination. First, Sgs1 is assumed to act downstream of mismatch recognition by Msh2 to unwind strand-exchange intermediates containing a high density of mismatches (Golfarb and Alani, 2005; Sugawara et al., 2004; Spell and Jinks-Robertson, 2004; Chakraborty and Alani, 2016). It is also possible that Sgs1 possesses Msh2-independent anti-recombination activity. Second, *SGS1* mutants have an increased number of cytologically visible connections between homologs, which could help to stabilize interactions between diverged chromosomes (Rockmill et al. 2003). Finally, Sgs1 also limits crossing over by facilitating recombinational repair without an associated exchange of chromosome arms (non-crossovers outcome; Bizard and Hickson, 2014). When *SGS1* expression is suppressed during meiosis, recombination intermediates are processed by structure-selective endonucleases to yield higher levels of crossovers (Oh et al, 2007; Zakaryevich et al. 2012; De Muyt et al. 2012; Rockmill et al. 2003). All of these factors may contribute to the increased rate of crossing over observed in the *SGS1* repression mutants.

### Genetic incompatibility as a barrier between species

In other organisms, hybrid sterility is shown to be caused by incompatibility between allele(s) from one species at one or more loci and allele(s) from the other species at one or more distinct loci (ex: Long et al., 2008; Mihola et al., 2009; Ting et al., 1998). BDMIs are expected to evolve quite readily when populations are isolated because new alleles, compatible with the genomes they evolve in, can spread by natural selection within their population, and are only costly if hybrids are formed with another isolated diverging population, combining new alleles that have not been together before. When experimental *S. cerevisiae* populations are evolved in divergent laboratory environments, hybrids between them have lower mitotic fitness in either environment (Dettman et al., 2007). Similar results are found when natural *S. cerevisiae* isolates and crosses between them are grown in a range of different laboratory conditions (Hou et al., 2015). These results suggest that genetic incompatibilities affecting F1 hybrid mitotic fitness occur readily within a species.

However F1 hybrids between *S. paradoxus* and *S. cerevisiae* do not show such incompatibilities in growth. On the contrary, they tend to show enhanced viability or “hybrid vigour” (Bernardes et al., 2017), so BDMIs for mitotic growth do not appear to be a major part of the species barrier between these well-established species. What of BDMIs affecting meiosis? Dettman et al. (2007) also report that their experimental hybrids show a relative reduction in “meiotic efficiency”, that is the proportion of diploids that enter meiosis when starved, but this is more a change of life history strategy than an intrinsically deleterious trait as presumably both unsporulated and sporulated cells remained viable. Well-defined mitochodrial-nuclear incompatibilites among *S. cerevisiae, S. paradoxus*, and *S. bayanus* can cause hybrids to lose the ability to respire, preventing entry into meiosis altogether (Lee et al., 2008; Chou et al., 2010). Such mito-nuclear incompatibilities may well reflect divergent adaptation of these different species (also see Xu and He, 2011).

Given the apparent ease with which incompatibilities affecting other parts of the yeast life cycle can evolve, it is surprising that BDMIs causing hybrid gamete inviability have not been detected to date (Xu and He, 2011; Kao et al., 2010). The largest and most direct attempt to identify BDMIs between nuclear genomes in yeast was conducted by Kao et al. (2010). They concluded that there were no “simple” BDMIs between the nuclear genomes of *S. cerevisiae* and *S. paradoxus*, where a simple BDMI is one that kills a certain hybrid genotype. They found several pairs of segments with distributions that were statistically significantly different than what would be expected by chance, but they attributed these to more complex interactions, likely involving multiple loci with weak effects. They found some evidence of three-way interactions, but lacked confidence due to limited statistical power. Li et al.’s simulation study explicitly investigated whether previous attempts at mapping BDMIs in yeast were adequate to conclude that they did not exist (Li et al., 2013). They found that BDMIs with incomplete penetrance (those do not kill all gametes of the incompatible genotype) would not be detected in Kao et al. (2010)’s study due to the limited sample size. Moreover, higher order interactions (involving three or more loci) would behave the same as incompletely penetrant two-way interactions and thus, given a sufficient sample size, could be detected statistically in the same way. They recommend using OR instead of ChiSq because it has the advantage of differentiating between differences due to over-representation and under-representation of a genotype relative to expectations.

We were able to build on the work of Kao et al. (2010) and Li et al. (2013) to successfully detect pairwise BDMIs. Using our *P*_*CLB2*_*-MSH2 P*_*CLB2*_*-SGS1* double mutant strains, we were able to restore recombination to an average of 37.9 cross-overs per meiosis (or 18.9 per spore). This was an improvement over Kao et al.’s deletion mutant of *MSH2*, which had an average of 17.8 recombination events per strain (Kao et al., 2010). They were also forced to exclude aneuploid chromosomes from most of their analysis, thus greatly decreasing the effective sample size (Kao et al., 2010). By obtaining complete tetrads, we avoided this problem. Moreover, we also reduced potential genotyping errors because each recombination event is supported in two separate, reciprocal samples. As well as these improvements, by using OR instead of ChiSq, as recommended by Li et al. (2013), we could focus solely on the case in which there is a depletion of hybrid types (OR higher than expected).

Using these improved methods, we found six major regions of the genome that appear to define four putative two-locus BDMIs (Figure 2). These regions were found on only four chromosomes (chr IX, X, XII and XV). Many genes map to these regions, and fine-scale mapping will be necessary to determine the causative loci. Among the known interacting genes in BioGRID, there are none identified between genes found in regions A and E or B and D (Oughtred et al., 2018). Among the genes in regions C and E, there is one known interaction; a negative genetic interaction between *IMA2* (an isomaltase) and *CDC6* (an essential protein required for DNA replication), which was found in a large-scale genetic interaction study (Costanzo et al., 2016). Regions D and F harbour many known interacting genes, but this is unsurprising because together they encompass the largest number of genes. Despite no known interactions between regions A and E or regions B and D, there are some good candidates based on similar proteins. For example, in region E, gene *YOL159C-A*, encoding a protein of unknown function, interacts positively with *COA4*, encoding a protein involved in the organization of cytochrome c oxidase (Cherry et al., 2012; Costanzo et al., 2016). *COA4* is not found in region A but *COA1*, which is also required for assembly of the cytochrome c oxidase complex, is. Additionally, *CSS3*, another protein of unknown function in region E, interacts negatively with *MAL12*, a maltase that hydrolyzes sucrose (Cherry et al., 2012; Costanzo et al., 2016). *MAL12* is not found in region A but *SUC2*, a sucrose hydrolyzing enzyme, is. In region B, there is only one gene, *CBR1*, a cytochrome b reductase. It has physical interactions with *GSC2*, a synthase involved in the formation of the inner layer of the spore wall (Cherry et al., 2012; Krogan et al., 2006) similar to *SPO75* found in region D, which is required for spore wall formation. It also interacts physically with *ORC1*, the largest subunit of the origin recognition complex (Cherry et al., 2012; Müller et al., 2010), and *ORC3*, another subunit of the origin recognition complex, is found in region D. *CBR1* also interacts negatively genetically with both *MDM36*, a mitochondrial protein which is proposed to have involvement in the formation of Dnm1p-containing cortical anchor complexes that promote mitochondrial fission, where *DNM1* is found in region D, and *WHI2*, a binding partner of Psr2p required for full activation of STRE-mediated gene expression, where *PSR2* is found in region D (Cherry et al., 2012; Costanzo et al., 2016). Whilst these incompatible regions contain interesting candidate genes, finer scale recombination mapping followed by candidate allele replacement and sensitive fertility assays will be necessary to determine the underlying molecular genetics of these BDMIs.

## Conclusion

Suppressing the expression of just two genes, *SGS1* and *MSH2*, rescues the fertility of normally sterile yeast hybrids and allows the recovery of recombinant euploid hybrid gametes, permitting the detection of nuclear-nuclear BDMIs in yeast for the first time. Whilst these incompatibilities comprise only a small part of the reproductive barrier between the parent species, with the vast majority coming from anti-recombination between the diverged genomes, they may have been important in the early stages of speciation. These results not only bring the power of yeast genetics to bear on the genetics of post zygotic reproductive isolation, they also enable crossing between highly diverged species, potentially allowing other interesting traits to be mapped or producing recombinant hybrids with novel commercial or research uses.

## Materials and Methods

### Strains

We used as a template a previously constructed *S. cerevisiae* strain NHY 2039, in which the promotor of *SGS1* had been replaced by the *CLB2* promotor (Oh et al., 2008) using the pFA6a-KANMX6-pCLB2-3HA created by Lee and Amon (2003). We amplified the *CLB2* promotor and the *KANMX4* drug resistance marker out of NHY2039 (i.e. YDG832) using primer pairs (see Supplementary File 5) that allowed us to transform it in place of the natural promoters of *MSH2* and *SGS1* in both *S. cerevisiae* (W303 background) and *S. paradoxus* (N17 background). The resulting *S. cerevisiae* and *S. paradoxus* haploid strains YDG968 and YDG969 (see Table 1, Supplementary File 1 and Supplementary File 2 for details) were crossed together producing an F1 hybrid diploid YDG982 in which both homologous copies of both *SGS1* and *MSH2* were under the control of the *CLB2* promotor, suppressing the expression of these genes during meiosis. To obtain a non-hybrid, double-mutant (i.e. *P*_*CLB2*_*-MSH2, P*_*CLB2*_*-SGS1*) control strain under the *S. paradoxus* background, we crossed haploid strains YDG967 and YDG968. Next, we crossed YDG969 and YDG970 strains to obtain a similar non-hybrid, double mutant (i.e. *P*_*CLB2*_*-MSH2, P*_*CLB2*_*-SGS1*) control strain for the *S. cerevisiae* background. Finally, to obtain a wild-type hybrid control strain (i.e. without CLB2 promoter replacement), we crossed haploid strains YDG391 (*S. paradoxus*) and YDG542 (*S. cerevisiae*), and selected for diploid clones (to form YDG853).

### Fertility

We induced meiosis and sporulation by incubating the hybrid diploid (YDG982) in 3 ml KAc (2% potassium acetate sporulation media) for four days in room temperature with vigorous shaking. To digest the ascus walls of the hybrid ascospores, we incubated them in 1unit (per 10 μl) zymolyase (Zymo Research EU, Freiburg, Germany) for 30 minutes. After enzymatic digestion of the ascus walls, we placed the four spores of each tetrad onto YEPD (2% glucose, 1% yeast extract, 2% peptone, 2% agar) plates using an MSM400 tetrad dissection microscope (Singer Instruments, Watchet, UK). Plates containing dissected tetrads were incubated at 30 °C before examining them for visible colonies founded by germinating spores.

We defined fertility as the proportion of viable gametes, i.e. the number of spores that germinated and formed colonies visible to the naked eye after 2 days, divided by the total number of spores that were dissected. For the hybrid crosses, we dissected a large number of spores (≧ 880, see Source Data 1). This was necessary for the hybrid crosses because they were known to have low gamete viability. For the non-hybrid crosses, we only dissected 384-400 spores (Source Data 1). Because the non-hybrid crosses had much higher rates of gamete viability than the hybrid crosses, dissecting a lower number of spores was sufficient to obtain a good estimate of their true fertility. Only technical replicates (repeated meioses of the same original diploid strain) were performed and were all considered to be part of a single sample.

### Sequencing and genotyping

To ensure that the hybrid gametes we sequenced were euploid, we only genotyped gametes from tetrads that contained four viable spores. Even with the observed 70-fold increase in hybrid gamete viability, only 5% of the tetrads contained four viable spores. In order to maximize useable data from a single lane of sequencing, we limited our sample size to 94 tetrads. Again, repeated meioses of a single diploid strain were performed and were all considered to be part of a single sample. We extracted DNA from all 376 colonies from 94 tetrads (in addition to two non-hybrid control tetrads) using MasterPure ™ Yeast DNA Purification Kit (Epicentre, Biozyme Biotech, Oldendorf, Germany). To prepare the samples for sequencing, we used double digestion based RAD-tag library preparation method (Etter et al., 2011; Hohenlohe et al., 2010; Peterson et al., 2012). We digested 50 ng of DNA from each colony using restriction enzymes *Csp*6I an*d Pst*I and ligated adapters (adapterX_TagY_fq and adapterX_TagY_rv) in the same reaction at 37 °C for two hours. We cleaned up the excess adapters, enzymes, and fragments smaller than 300bp by using Ampure beads at a 1:1 ratio. Next we mixed Phusion Hot Start II High-Fidelity DNA Polymerase (2U/µl), adding P5 and P7 primers at 10 mM concentration, dNTPs (2mM per dNTP), and 5X Phusion HF Buffer to amplify the target regions (Acinas et al., 2005; Etter et al., 2011). 30 µl PCR mixtures were amplified as an initial 98 °C incubation for 30s, followed by 25 cycles of 98 °C for 10s, 68 °C for 15s, 72 °C for 30s, and then a final extension at 72 °C for 5 mins. To sequence the tagged samples, we mixed all tagged samples in one pool. All samples were multiplexed using combinations of 24 unique barcodes therefore reads from a single sequencing reaction would have unique reverse and forward tags that will help us to distinguish all samples after obtaining the pool of MiSeq reads. We used MiSeq platform to obtain 300 bp paired-end reads. Raw sequence data is available from Dryad (Bozdag et al., 2019).

To map the reads, we assembled two simplified co-linear reference genomes consisting of the coding DNA only from the set of open reading frames shared between *S. cerevisiae* and *S. paradoxus*, removing open reading frames that were present in one species but not in another or which were not co-linear (based on SGRP sequencing added 10/10/08; Liti et al., 2009; Bergström et al., 2014). We mapped reads to these reference genomes using NGK. At this point, we excluded 10 tetrads due to poor sequencing coverage and quality, leaving us with 336 samples from 84 tetrads. We assigned ORFs to one or other species using two simplifying assumptions: that no non-Mendelian segregation occurred and that recombination occurred only in intergenic regions. Thus, if reads in all four spores of a tetrad contained reads mapping to a given ORF of one or the other or both species, the two spores with the highest proportion of reads mapping to one species ORF would have it assigned to that species and the other two would have the ORF assigned to the other species. If the four copies of an ORF within a tetrad did not all contain reads mapping to either or both species then the ORF would be assigned to the same species as the neighbouring ORF. These genotyping rules produced a recombination map (see Supplementary File 6 for an example, full data available in Bozdag et al., 2019) of the four spores within each tetrad at ORF-level resolution, with no gain or loss of genetic material (i.e. no gene-conversion).

## Analysis

We divided the chromosomes into 1208 segments defined by the recombination breakpoints observed in all 84 tetrads (Supplementary File 3), reasoning that we could not resolve a locus smaller that the closest crossovers flanking it. Each segment was tested against each other segment, excluding pairs found on the same chromosome, similar to the method described by Li et al. (2013) (exact method can be found in Source Code 1). Within chromosome pairs were not tested because physical linkage would skew the numbers towards parental combinations, the same effect that we expect to see due to incompatibility between loci, thus making the results difficult to interpret. Following the procedure of Li et al. (2013), the odds ratio (OR) was calculated for each pair of segments by dividing the product of the numbers of the two parental genotypes observed by that of the two hybrid types:

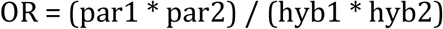

Pairs of segments on the same chromosome were excluded because physical linkage would skew the numbers towards parental combinations, the same effect that we expect to see due to incompatibility between loci, thus making the results difficult to interpret.

To control for type I error, we produced a null distribution against which to test our observed ORs (Source Code 1). First, a new set of 84 tetrads was simulated by pulling each chromosome for each tetrad from the pool of chromosomes of that type (with replacement). This is similar to, but not exactly the same as, the method used by Li et al. (2013). They, in contrast, shuffle the chromosomes without replacement instead of sampling with replacement, and they do so among random spores instead of whole tetrads. ORs for the segment pairs were then calculated as before but on the simulated set of chromosomes. This process was repeated 99 more times, and the top OR from each set of simulated tetrads was recorded. The 5th largest OR from this set of 100 top ORs was chosen as the critical value from which to judge significance. All observed pairs with a higher OR than this critical value (3.4131) were deemed statistically significant (Supplementary File 4 and Source Data 2). For our sample size of 84 tetrads, this represents a 1.85-fold over-representation of parental combinations (218 parental combinations vs. 118 hybrid combinations for that pair of alleles).

The significant pairs of segments were then grouped into blocks comprising neighbouring pairs. Instead of looking at blocks of seven markers, as was done in Li et al. (2013), we decided that if two adjacent segments both had a significant interaction with the same segment in another chromosome, they were part of the same block. In one case, one significant interaction was between a non-adjacent segment and a segment that interacted with many other nearby segments. Because this interaction was so close to the others (within nine segments), we arbitrarily decided to treat it as part of the same interaction (Source Data 2, row 37 as compared to surrounding rows). Similarly, region B was found to interact with a segment adjacent to those interacting with region F. In this case, we collapsed both interacting regions into one (region D), as we consider it most likely that the two regions are interacting with a single gene in the region.

## Acknowledgements

Numerous people have contributed to this long-running project over the years, and we apologise if we have neglected to name you. We are particularly grateful to colleagues at the Max Planck Institute for Evolutionary Biology, especially Arne Nolte, Gunda Dechow-Seligmann, and Elke Bustorf, who genotyped our hybrid strains. Mahesh Binzer-Panchal de-multiplexed the sequence reads. Krishna B. S. Swamy provided invaluable input on data analysis. Michael Scott at UCL generously helped us with code and simulations.

## Supplementary Material

Supplementary File 1: A schematic of how the strains used in this study were constructed. Supplementary File 2: Complete list of strains used in this study.

Supplementary File 3: Segments used for mapping BDMIs. Each segment is defined by observed recombination between genes and all start and end locations are based on the sequence of W303.

Supplementary File 4: All pairwise combinations of segments, the observed number of parental and hybrid combinations and the summary statistics.

Supplementary File 5: Primers used in the study.

Supplementary File 6: An example recombination map of a single tetrad. Gametes were genotyped by ORF into one of the two species, ensuring a 2:2 segregation of species identity at each ORF.

Source Data 1: Raw counts of dissected and viable spores for each strain assayed in this study. Source Data 2: Significant pairwise combinations of segments grouped by interacting regions.

Source Code 1: Code used to perform statistical analysis on gene interaction data, along with necessary input files.

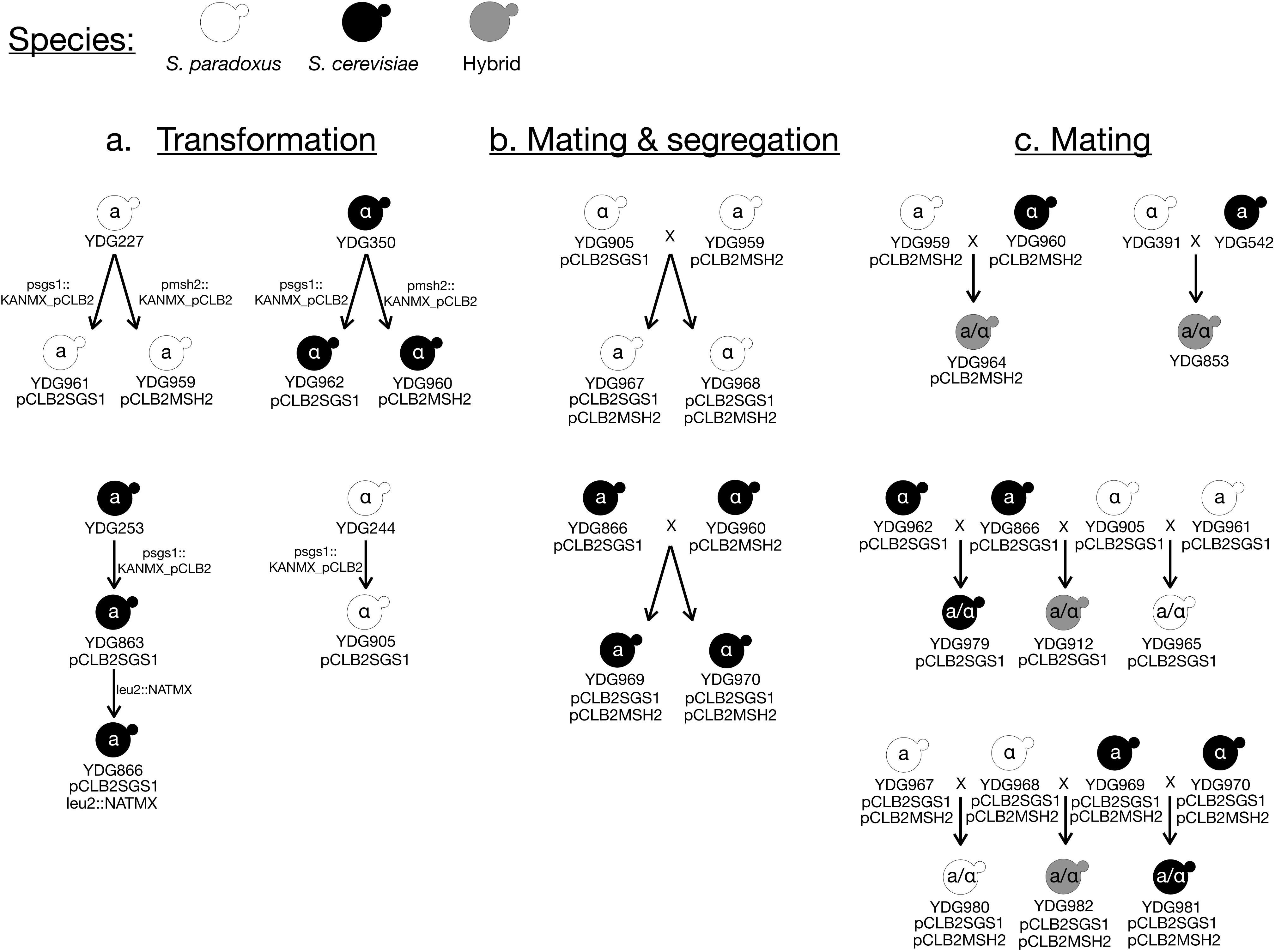

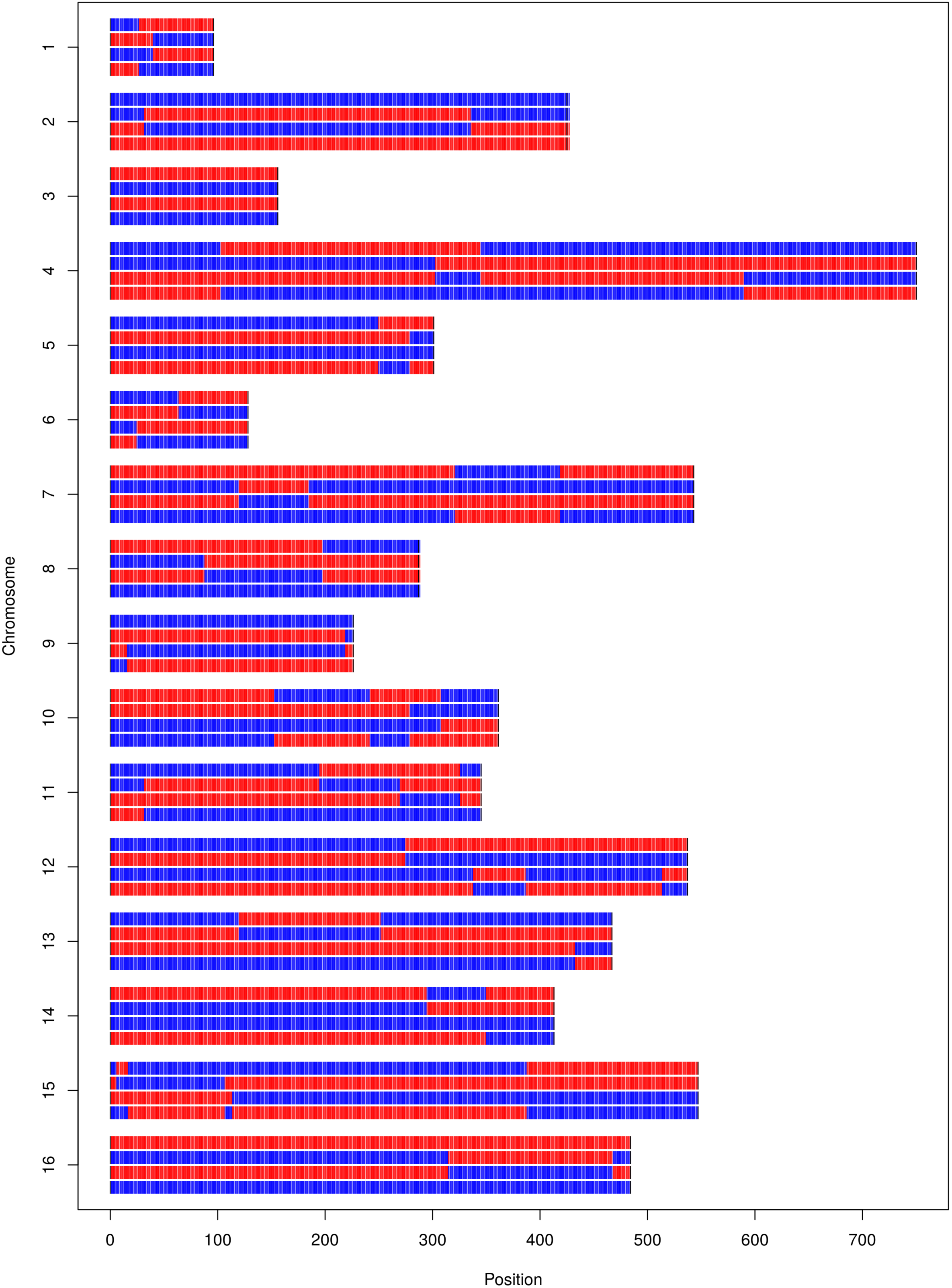

## References

Acinas, S. G., Sarma-Rupavtarm, R., Klepac-Ceraj, V., & Polz, M. F. (2005). PCR-induced sequence artifacts and bias: insights from comparison of two 16S rRNA clone libraries constructed from the same sample. Appl. Environ. Microbiol., 71(12), 8966–8969. doi: 10.1128/AEM.71.12.8966-8969.2005

Amin, A. D., Chaix, A. B., Mason, R. P., Badge, R. M., & Borts, R. H. (2010). The roles of the Saccharomyces cerevisiae RecQ helicase SGS1 in meiotic genome surveillance. PloS one, 5(11), e15380. https://doi.org/10.1371/journal.pone.0015380

Bergström, A., Simpson, J. T., Salinas, F., Barré, B., Parts, L., Zia, A., … & Warringer, J. (2014). A high-definition view of functional genetic variation from natural yeast genomes. Molecular biology and evolution, 31(4), 872–888. https://doi.org/10.1093/molbev/msu037

Bernardes, J. P., Stelkens, R. B., & Greig, D. (2017). Heterosis in hybrids within and between yeast species. Journal of Evolutionary Biology, 30(3), 538–548. https://doi.org/10.1111/jeb.13023

Bizard, A. H., & Hickson, I. D. (2014). The dissolution of double Holliday junctions. Cold Spring Harbor perspectives in biology, 6(7), a016477. doi: 10.1101/cshperspect.a016477

Botstein, D., & Fink, G. R. (2011). Yeast: an experimental organism for 21st Century biology. Genetics, 189(3), 695–704. https://doi.org/10.1534/genetics.111.130765

Boynton, P. J., Janzen, T., & Greig, D. (2018). Modeling the contributions of chromosome segregation errors and aneuploidy to Saccharomyces hybrid sterility. Yeast, 35(1), 85–98. https://doi.org/10.1002/yea.3282

Bozdag, G. O., Ono, J., Denton, J. A., Karakoc, E., Hunter, N., Leu, J.-Y., and Greig, D. (2019). Data from: Engineering recombination between diverged yeast species reveals speciation genes. Dryad Digital Repository. doi:10.5061/dryad.bk02240

Chakraborty, U., & Alani, E. (2016). Understanding how mismatch repair proteins participate in the repair/anti-recombination decision. FEMS yeast research, 16(6), fow071. https://doi.org/10.1093/femsyr/fow071

Charron, G., Leducq, J. B., & Landry, C. R. (2014). Chromosomal variation segregates within incipient species and correlates with reproductive isolation. Molecular ecology, 23(17), 4362–4372. https://doi.org/10.1111/mec.12864

Cherry, J. M., Hong, E. L., Amundsen, C., Balakrishnan, R., Binkley, G., Chan, E. T., … & Fisk, D. G. (2012). Saccharomyces Genome Database: the genomics resource of budding yeast. Nucleic acids research, 40(D1), D700–D705. https://doi.org/10.1093/nar/gkr1029

Chou, J. Y., Hung, Y. S., Lin, K. H., Lee, H. Y., & Leu, J. Y. (2010). Multiple molecular mechanisms cause reproductive isolation between three yeast species. PLoS biology, 8(7), e1000432. https://doi.org/10.1371/journal.pbio.1000432

Costanzo, M., VanderSluis, B., Koch, E. N., Baryshnikova, A., Pons, C., Tan, G., et al. (2016). A global genetic interaction network maps a wiring diagram of cellular function. Science, 353(6306), aaf1420–aaf1420. doi: 10.1126/science.aaf1420

Coyne, J. A., & Orr, H. A. (2004). Speciation. Sinauer. Sunderland, MA.

De Muyt, A., Jessop, L., Kolar, E., Sourirajan, A., Chen, J., Dayani, Y., & Lichten, M. (2012). BLM helicase ortholog Sgs1 is a central regulator of meiotic recombination intermediate metabolism. Molecular cell, 46(1), 43–53. https://doi.org/10.1016/j.molcel.2012.02.020

Dettman, J. R., Sirjusingh, C., Kohn, L. M., & Anderson, J. B. (2007). Incipient speciation by divergent adaptation and antagonistic epistasis in yeast. Nature, 447(7144), 585–588. https://doi.org/10.1038/nature05856

Etter, P. D., Bassham, S., Hohenlohe, P. A., Johnson, E. A., & Cresko, W. A. (2011). SNP Discovery and Genotyping for Evolutionary Genetics Using RAD Sequencing. Methods in molecular biology (Clifton, NJ), 772, 157. https://doi.org/10.1007/978-1-61779-228-1_9

Fischer, G., James, S. A., Roberts, I. N., Oliver, S. G., & Louis, E. J. (2000). Chromosomal evolution in Saccharomyces. Nature, 405(6785), 451. https://doi.org/10.1038/35013058

Goldfarb, T., & Alani, E. (2005). Distinct roles for the Saccharomyces cerevisiae mismatch repair proteins in heteroduplex rejection, mismatch repair and nonhomologous tail removal. Genetics, 169(2), 563–574. https://doi.org/10.1534/genetics.104.035204

Grandin, N., & Reed, S. I. (1993). Differential function and expression of Saccharomyces cerevisiae B-type cyclins in mitosis and meiosis. Molecular and cellular biology, 13(4), 2113–2125. doi: 10.1128/MCB.13.4.2113

Greig, D. (2007). A screen for recessive speciation genes expressed in the gametes of F1 hybrid yeast. PLoS genetics, 3(2), e21. https://doi.org/10.1371/journal.pgen.0030021

Greig, D., Travisano, M., Louis, E. J., & Borts, R. H. (2003). A role for the mismatch repair system during incipient speciation in Saccharomyces. Journal of evolutionary biology, 16(3), 429–437. https://doi.org/10.1046/j.1420-9101.2003.00546.x

Hohenlohe, P. A., Bassham, S., Etter, P. D., Stiffler, N., Johnson, E. A., & Cresko, W. A. (2010). Population genomics of parallel adaptation in threespine stickleback using sequenced RAD tags. PLoS genetics, 6(2), e1000862. https://doi.org/10.1371/journal.pgen.1000862

Hou, J., Friedrich, A., Gounot, J.-S., & Schacherer, J. (2015). Comprehensive survey of condition-specific reproductive isolation reveals genetic incompatibility in yeast. Nature Communications, 6, 7214. https://doi.org/10.1038/ncomms8214

Hunter, N., Chambers, S. R., Louis, E. J., & Borts, R. H. (1996). The mismatch repair system contributes to meiotic sterility in an interspecific yeast hybrid. The EMBO journal, 15(7), 1726–1733. https://doi.org/10.1002/j.1460-2075.1996.tb00518.x

Kao, K. C., Schwartz, K., & Sherlock, G. (2010). A genome-wide analysis reveals no nuclear Dobzhansky-Muller pairs of determinants of speciation between S. cerevisiae and S. paradoxus, but suggests more complex incompatibilities. PLoS genetics, 6(7), e1001038. https://doi.org/10.1371/journal.pgen.1001038

Kellis, M., Patterson, N., Endrizzi, M., Birren, B., & Lander, E. S. (2003). Sequencing and comparison of yeast species to identify genes and regulatory elements. Nature, 423(6937), 241. https://doi.org/10.1038/nature01644

Krogan, N. J., Cagney, G., Yu, H., Zhong, G., Guo, X., Ignatchenko, A., … & Punna, T. (2006). Global landscape of protein complexes in the yeast Saccharomyces cerevisiae. Nature, 440(7084), 637. https://doi.org/10.1038/nature04670

Lee, B. H., & Amon, A. (2003). Role of Polo-like kinase CDC5 in programming meiosis I chromosome segregation. Science, 300(5618), 482–486. doi: 10.1126/science.1081846

Lee, H. Y., Chou, J. Y., Cheong, L., Chang, N. H., Yang, S. Y., & Leu, J. Y. (2008). Incompatibility of nuclear and mitochondrial genomes causes hybrid sterility between two yeast species. Cell, 135(6), 1065–1073. https://doi.org/10.1016/j.cell.2008.10.047

Li, C., Wang, Z., & Zhang, J. (2013). Toward Genome-Wide Identification of Bateson – Dobzhansky–Muller Incompatibilities in Yeast: A Simulation Study. Genome biology and evolution, 5(7), 1261–1272. https://doi.org/10.1093/gbe/evt091

Liti, G., Carter, D. M., Moses, A. M., Warringer, J., Parts, L., James, S. A., … & Tsai, I. J. (2009). Population genomics of domestic and wild yeasts. Nature, 458(7236), 337. https://doi.org/10.1038/nature07743

Long, Y., Zhao, L., Niu, B., Su, J., Wu, H., Chen, Y., … & Xia, J. (2008). Hybrid male sterility in rice controlled by interaction between divergent alleles of two adjacent genes. Proceedings of the National Academy of Sciences, 105(48), 18871–18876. https://doi.org/10.1073/pnas.0810108105

Maheshwari, S., & Barbash, D. A. (2011). The genetics of hybrid incompatibilities. Annual review of genetics, 45, 331–355. https://doi.org/10.1146/annurev-genet-110410-132514

Martini, E., Diaz, R. L., Hunter, N., & Keeney, S. (2006). Crossover homeostasis in yeast meiosis. Cell, 126(2), 285–295. https://doi.org/10.1016/j.cell.2006.05.044

Mihola, O., Trachtulec, Z., Vlcek, C., Schimenti, J. C., & Forejt, J. (2009). A mouse speciation gene encodes a meiotic histone H3 methyltransferase. Science, 323(5912), 373–375. doi: 10.1126/science.1163601

Müller, P., Park, S., Shor, E., Huebert, D. J., Warren, C. L., Ansari, A. Z., … & Fox, C. A. (2010). The conserved bromo-adjacent homology domain of yeast Orc1 functions in the selection of DNA replication origins within chromatin. Genes & development, 24(13), 1418–1433. doi: 10.1101/gad.1906410

Nosil, P., & Schluter, D. (2011). The genes underlying the process of speciation. Trends in Ecology & Evolution, 26(4), 160–167. https://doi.org/10.1016/j.tree.2011.01.001

Oh, S. D., Lao, J. P., Hwang, P. Y. H., Taylor, A. F., Smith, G. R., & Hunter, N. (2007). BLM ortholog, Sgs1, prevents aberrant crossing-over by suppressing formation of multichromatid joint molecules. Cell, 130(2), 259–272. https://doi.org/10.1016/j.cell.2007.05.035

Oh, S. D., Lao, J. P., Taylor, A. F., Smith, G. R., & Hunter, N. (2008). RecQ helicase, Sgs1, and XPF family endonuclease, Mus81-Mms4, resolve aberrant joint molecules during meiotic recombination. Molecular cell, 31(3), 324–336. https://doi.org/10.1016/j.molcel.2008.07.006

Orr, H. A. (1996). Dobzhansky, Bateson, and the genetics of speciation. Genetics, 144(4), 1331.

Oughtred, R., Stark, C., Breitkreutz, B. J., Rust, J., Boucher, L., Chang, C., … & Zhang, F. (2018). The BioGRID interaction database: 2019 update. Nucleic acids research, 47(D1), D529–D541. https://doi.org/10.1093/nar/gky1079

Peterson, B. K., Weber, J. N., Kay, E. H., Fisher, H. S., & Hoekstra, H. E. (2012). Double digest RADseq: an inexpensive method for de novo SNP discovery and genotyping in model and non-model species. PloS one, 7(5), e37135. https://doi.org/10.1371/journal.pone.0037135

Presgraves, D. C. (2010). The molecular evolutionary basis of species formation. Nature Reviews Genetics, 11(3), 175. https://doi.org/10.1038/nrg2718

Rieseberg, L. H., & Blackman, B. K. (2010). Speciation genes in plants. Annals of Botany, 106(3), 439–455. https://doi.org/10.1093/aob/mcq126

Rieseberg, L. H., & Willis, J. H. (2007). Plant speciation. Science, 317(5840), 910–914. doi: 10.1126/science.1137729

Rockmill, B., Fung, J. C., Branda, S. S., & Roeder, G. S. (2003). The Sgs1 helicase regulates chromosome synapsis and meiotic crossing over. Current Biology, 13(22), 1954–1962. https://doi.org/10.1016/j.cub.2003.10.059

Rogers, D. W., McConnell, E., Ono, J., & Greig, D. (2018). Spore-autonomous fluorescent protein expression identifies meiotic chromosome mis-segregation as the principal cause of hybrid sterility in yeast. PLoS biology, 16(11), e2005066. https://doi.org/10.1371/journal.pbio.2005066

Spell, R. M., & Jinks-Robertson, S. (2004). Examination of the roles of Sgs1 and Srs2 helicases in the enforcement of recombination fidelity in Saccharomyces cerevisiae. Genetics, 168(4), 1855–1865. https://doi.org/10.1534/genetics.104.032771

Sugawara, N., Goldfarb, T., Studamire, B., Alani, E., & Haber, J. E. (2004). Heteroduplex rejection during single-strand annealing requires Sgs1 helicase and mismatch repair proteins Msh2 and Msh6 but not Pms1. Proceedings of the National Academy of Sciences, 101(25), 9315–9320. https://doi.org/10.1073/pnas.0305749101

Ting, C. T., Tsaur, S. C., Wu, M. L., & Wu, C. I. (1998). A rapidly evolving homeobox at the site of a hybrid sterility gene. Science, 282(5393), 1501–1504. doi: 10.1126/science.282.5393.1501

Xu, M., & He, X. (2011). Genetic incompatibility dampens hybrid fertility more than hybrid viability: yeast as a case study. PLoS One, 6(4), e18341. https://doi.org/10.1371/journal.pone.0018341

Zakharyevich, K., Tang, S., Ma, Y., & Hunter, N. (2012). Delineation of joint molecule resolution pathways in meiosis identifies a crossover-specific resolvase. Cell, 149(2), 334–347. https://doi.org/10.1016/j.cell.2012.03.023

